# Profiling Lysosomal and Mitochondrial Dysfunction in Neurodegenerative Diseases Using Human Fibroblasts for Translational Therapeutic Screening

**DOI:** 10.1101/2025.06.24.661057

**Authors:** Diana Moreira Leite, Rhianna Lenham, Philippa Malko, Stuart Thomson, Tim Phillips, Tatiana Rosado Rosenstock, Naheed Rohman Mirza

## Abstract

Lysosomal dysfunction and mitochondrial health are intricately connected, playing essential roles in cellular homeostasis. Lysosomes are acidic membrane-bound organelles responsible for degrading and recycling cellular waste, while mitochondria generate the energy required for cellular functions. Growing evidence implicates roles for lysosomal and mitochondrial dysfunction in neurodegenerative diseases, including Alzheimer’s and Parkinson’s disease. With novel therapeutics targeting both the lysosomal and mitochondrial functions, robust assays for compound screening are becoming critical to evaluate modulation of both organelles in disease-relevant cellular models. Here, we investigated human fibroblasts derived from healthy donors, as well as patients with Alzheimer’s and Parkinson’s disease, to assess their capacity to model key aspects of lysosomal and mitochondrial dysfunction. Lysosomal function was evaluated using various assays, including quantification of lysosomal proteins (TMEM175 and LAMP1), LysoTracker™ staining, measurement of lysosomal pH, and lysosomal enzymatic activity. Autophagic flux was assessed by measuring p62 levels as a marker of autophagy. Mitochondrial function was investigated by measuring mitochondrial calcium levels, membrane potential, oxidative stress, and mitochondrial content using MitoTracker™. To explore the potential of using human fibroblasts *for in vitro* compound screening, we validated these assays in a 384-well high-throughput format using compounds such as chloroquine and ammonium chloride. Our findings demonstrate that human fibroblasts faithfully recapitulate lysosomal and mitochondrial dysfunctions characteristic of neurodegenerative diseases. Moreover, the use of robust assays positions these cells as a valuable platform for high-throughput screening to identify novel therapeutics targeting lysosomal and mitochondrial pathways.

## Introduction

Lysosomes are acidic membrane-bound organelles that serve as a degradation and recycling centre of the cell, while also functioning as a signalling hub and participating in multiple key biological processes, including membrane repair, homeostasis, and immune response ^1,2^. Mitochondria are highly dynamic organelles critical for homeostasis maintenance and stress response. As the sites for biochemical processes, such as adenosine triphosphate (ATP) production, fatty acids synthesis, generation of intracellular reactive oxygen species (ROS), oxidative phosphorylation, and calcium homeostasis, mitochondria are also a hub of cellular metabolism ^3^. Lysosomal and mitochondrial functions are intricately connected ^4–6^. Evidence has revealed that these organelles communicate directly via membrane contact sites, facilitating the exchange of lipids, calcium, and other signalling molecules ^6^. This interaction is crucial for processes, such as mitophagy, in which damaged mitochondria are identified and degraded by lysosomes to prevent cellular stress and maintain metabolic balance. Recent growing evidence implicates lysosomal and mitochondrial dysfunctions in the progression of diseases, such as Alzheimer’s disease (AD) and Parkinson’s disease (PD) ^7–11^. AD and PD are the two most prevalent age-related neurodegenerative disorders, each characterised by distinct clinical and pathological features, but sharing overlapping molecular mechanisms, particularly involving mitochondrial and lysosomal dysfunction.

AD primarily presents with progressive cognitive decline and memory impairment ^12^. Its pathological hallmarks include extracellular accumulation of amyloid-β (Aβ) plaques and intracellular aggregates of hyperphosphorylated Tau ^12^. Emerging evidence has linked these proteinopathies to disruptions in mitochondrial function, such as impaired electron transport chain activity, elevated oxidative stress, and dysregulated mitochondria dynamics ^13–15^, as well as lysosomal ^16^. Dysfunctional lysosomal enzymatic activity, impaired clearance mechanisms, and accumulation of autophagic and lysosomal vesicles adjacent Aβ deposits suggest that compromised degradative pathways possess a key role in disease progression ^16^. Genetic studies also uncovered associations between AD and lysosomal genes, indicating a broader role for endolysosomal trafficking in disease aetiology ^17^.

PD, while primarily known for its motor symptoms (bradykinesia, tremor, rigidity, and postural instability) ^18^, is driven by the progressive loss of dopaminergic neurons in the substantia nigra and the accumulation of α-synuclein (α-syn) in Lewy bodies. Similar to AD, mitochondrial and lysosomal impairments are central to PD pathology ^19–21^. Familial forms of PD are associated with at least six mutated genes, including SNCA, PINK1, PRKN, DJ-1 and LRRK2, which regulate mitochondrial homeostasis and autophagy. Moreover, sporadic and familial PD appear to exhibit mitochondrial dysfunctions, such as disruption of the mitochondrial electron transport chain ^22^, deletions in mitochondrial DNA ^23^, as well as defects in regulation of mitochondrial dynamics ^24^. Mutations in various lysosomal genes (ATP13A2, TMEM175, CTSB, SCARB2, ATP6V0A1, GALC, GUSB and NEU1) have been implicated in monogenic and sporadic PD forms, further highlighting lysosomal vulnerability ^25,26^.

These shared pathological features highlight a growing consensus that mitochondrial dysfunction and lysosomal impairment are not isolated events, but rather interconnected contributors to neurodegeneration. Exploring the intersection of these pathways in AD and PD may unveil new therapeutic strategies to restore cellular homeostasis across various neurodegenerative disorders. Therefore, it is critical to establish robust and scalable assays to evaluate mitochondrial and lysosomal function in relevant cellular models. Human fibroblasts derived from patients represent a valuable platform for screening therapeutic modalities, as these cell models may retain specific mutations and dysfunctional pathways relevant to the disease ^27^. Additionally, fibroblasts from both healthy donors and affected individuals are more accessible, robust, and amenable to high-throughput screening (HTS) compared to other cellular models, such as primary rodent cultures or induced pluripotent stem cells ^28^.

In this study, we first explored whether fibroblasts from both AD and PD donors exhibit lysosomal, autophagy, and mitochondrial dysfunctions similar to those reported in these disorders. We then focused on PD fibroblasts to profile two compounds, including chloroquine (CQ) and ammonium chloride (AC), to validate lysosomal assays suitable for inclusion in drug discovery screening cascades. Ultimately, this approach offers new opportunities to accelerate the innovation of disease-modifying therapies for AD, PD, and related neurodegenerative disorders.

## Materials and Methods

### Cell Culture

Fibroblasts derived from healthy, familial AD, and idiopathic PD donors were obtained from the Coriell Cell Repository (**Table 1**). Donors were age- and sex-matched.

**Table 1.**
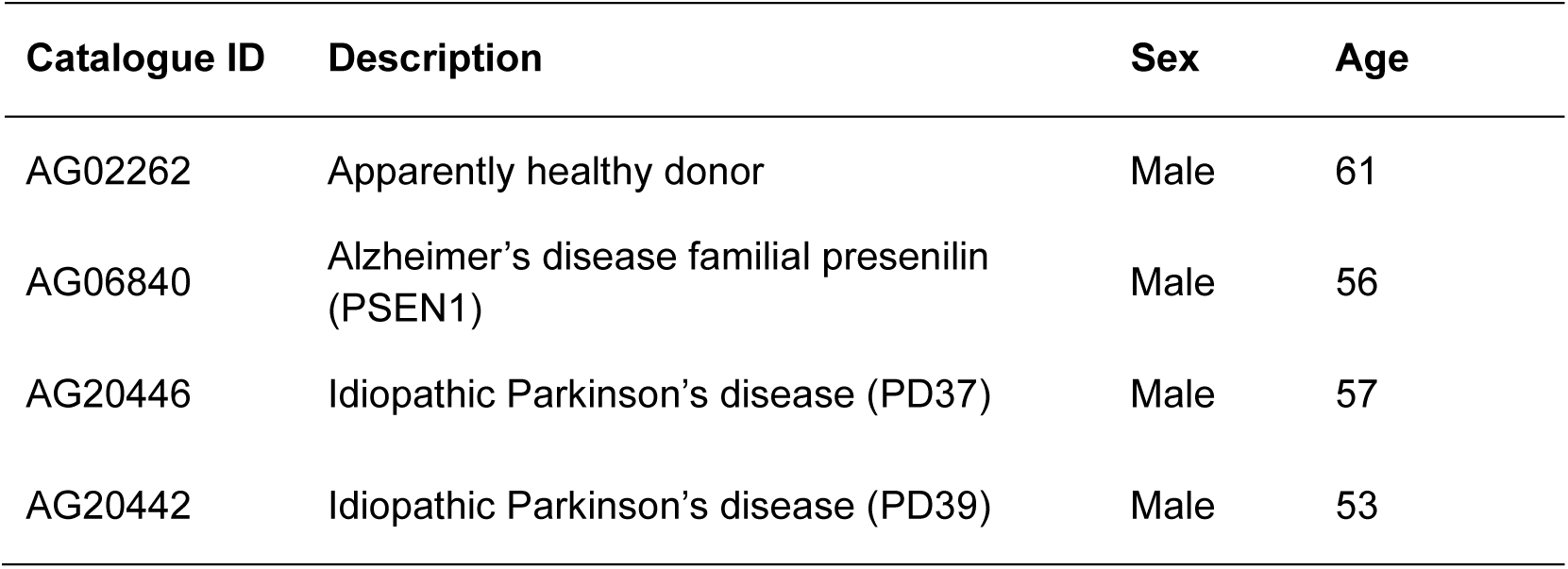
Details of human fibroblasts used in this study.

| Catalogue ID | Description | Sex | Age |
| --- | --- | --- | --- |
| AG02262 | Apparently healthy donor | Male | 61 |
| AG06840 | Alzheimer's disease familial presenilin (PSEN1) | Male | 56 |
| AG20446 | Idiopathic Parkinson's disease (PD37) | Male | 57 |
| AG20442 | Idiopathic Parkinson's disease (PD39) | Male | 53 |

Healthy and AD fibroblasts were maintained in Minimum Essential Medium (MEM, Sigma-Aldrich) supplemented with 15% (v/v) non-heat-inactivated foetal bovine serum (FBS, Sigma-Aldrich), 1% (v/v) non-essential amino acids (NEAA, Sigma-Aldrich) and 1% (v/v) penicillin/streptomycin (PS, Gibco). PD-derived fibroblasts were cultured in a mixture of MEM and Dulbecco’s Modified Eagle Medium (DMEM, Gibco) (1:1), 15% (v/v) non-heat inactivated FBS, 1% (v/v) NEAA and 1% (v/v) PS. This medium is referred to as complete medium. Cells were used at passage < 20 and routinely cultured as recommended by the manufacturer’s protocol.

### TMEM175, LAMP1 and p62 Immunofluorescence Assays

For TMEM175, LAMP1 and p62 immunofluorescence analysis, fibroblasts were seeded at 2,500 cells/well in a 384-well plate in 40 µL of complete media. To trigger autophagy and lysosomal activity, cells were maintained under serum starvation. For serum starvation, the media was replaced by serum-free media, and cells were incubated for 4 or 18 hours. At the end of the incubation period, fibroblasts were fixed with 4% (v/v) paraformaldehyde (PFA, Invitrogen), washed twice with phosphate-buffered saline (PBS), and then incubated with 10% (v/v) heat-inactivated FBS and 0.1% (v/v) Triton X-100 (Sigma-Aldrich) in PBS for 1 hour at 37 °C.

Primary antibodies were incubated overnight at 4 °C in 1% (v/v) FBS and 0.3% (v/v) Triton X-100 in PBS. Antibodies for TMEM175 (703805, ThermoFisher), LAMP1 (ab25630, Abcam), and p62 (ab109012, Abcam) were used at 1:1000, 1:500, and 1:2000, respectively. CF488 goat anti-rabbit and/or CF647 goat anti-mouse (BT20019 and BT20281, Cambridge Bioscience) secondary antibodies were added for 1 hour at 37 °C. Following PBS washing, samples were incubated with Hoechst (Thermo Scientific, 2 µM) for 10 minutes.

Fibroblasts were imaged on a high-content ImageXpress Micro Confocal instrument (Molecular Devices). The fluorescence intensity of TMEM175, LAMP1, and p62 was quantified using the Multi-Wavelength Cell Scoring Application Module for MetaXpress software (Molecular Devices) and normalised per cell number (nuclei staining). Four images were acquired per well (n=4 wells per condition, N=3 independent experiments).

### LysoTracker^TM^ Deep Red Assay

To investigate lysosome content, fibroblasts were seeded at 2,500 cells/well in a 384-well plate in 40 µL of complete media, and cells were serum starved for 18 hours. LysoTracker^TM^ Deep Red (L12492, Invitrogen) was used at a concentration of 10 nM and co-incubated with Hoechst (2 µM). Following an incubation for 30 minutes at 37 °C, cells were washed twice using PBS and imaged in Live Cell Imaging Media (Invitrogen) using ImageXpress Micro Confocal instrument.

LysoTracker^TM^ fluorescence intensity was measured using Multi-Wavelength Cell Scoring Application Module for MetaXpress software and normalised per cell number (nuclei staining). Four images were acquired per well (n=4 well per condition, N=3 independent experiments).

### LysoSensor^TM^ Yellow/Blue DND-160 Assay

To assess lysosomal pH changes, fibroblasts were seeded at 5,000 cells/well in a black 384-well plate and incubated overnight in complete media. LysoSensor^TM^ Yellow/Blue DND-160 (L7545, Invitrogen) was added at 20 µM, and incubated for 5 minutes at 37 °C. Cells were washed with PBS, and dual fluorescence signal was measured at λ_exc_ 329/λ_em_ 440 and λ_exc_ 384 /λ_em_ 540 using EnVision Plate Reader (Revvity). Yellow/Blue ratio, which indicates the levels of acidic lysosomes, was calculated by dividing the fluorescence values obtained from 384/540 nm by 329/440 nm. Three wells per condition (n=3 well per condition, N=3 independent experiments).

### Magic Red Cathepsin B Assay

To evaluate Cathepsin B activity, fibroblasts were seeded at 2,500 cells/well in a 384-well plate in 40 µL of complete media. For the serum starvation, the media was replaced by serum-free media, and cells were incubated for an additional 18 hours. Magic Red Cathepsin B substrate (ab270772, Abcam) was added (1:25 dilution in media) and incubated for 30 minutes at 37 °C. Nuclei staining was carried out by adding Hoechst (2 µM) to the Magic Red solution. Cells were washed with PBS and imaged in Live Cell Imaging Media using ImageXpress Micro Confocal instrument. Cathepsin B activity was quantified using Multi-Wavelength Cell Scoring Application Module for MetaXpress software to quantify total fluorescence and normalised per cell number (nuclei staining). Four images were acquired per well (n=4 well per condition, N=3 independent experiments).

### Mitochondrial Calcium, Membrane Potential and Oxidative Stress Assays

For all the mitochondrial assays, fibroblasts were seeded at 30,000 cells/well in a black 96-well plate in 200 µL of complete media. Measurements of mitochondrial calcium, membrane potential and reactive oxygen species (ROS) were performed using 10 µM Fluo-3 AM (F1242, Invitrogen), 250 nM tetramethyl rhodamine ethyl ester (TMRE, T669, Invitrogen) and 20 µM chloromethyl derivative of 2’, 7’-dichlorodihydrofluorescein diacetate (CM-H_2_DCFDA, C6827, Invitrogen), respectively ^29,30^. Cells were loaded with Fluo-3 AM in Krebs-Ringer solution supplemented with 1 mM calcium chloride and 1% (v/v) Pluronic F-27 and incubated for 1 hour at 37 °C. For membrane potential and ROS measurements, fibroblasts were either loaded with TMRE or CM-H_2_DCFDA in Krebs-Ringer solution supplemented with 1 mM calcium chloride and incubated for 1 hour or 30 minutes, respectively. Fluorescence was measured using a PHERAstar Microplate Reader (BMG Labtech) at 10 second intervals for a period of 5 minutes to obtain basal levels, and then for 5 minutes after addition of the mitochondrial uncoupler fluorocarbonyl cyanide phenylhydrazone (FCCP, C2920, Sigma-Aldrich) at 10 µM. Delta (Δ) values were calculated by subtracting the basal fluorescence (calculated from the average of the 10 readings prior to FCCP addition) from the post-FCCP fluorescence. Experiments were carried out in triplicate (n=3 wells per condition, N=4-5 independent experiments).

### MitoTracker^TM^ Deep Red Assay

Fibroblasts were seeded at 30,00 cells/well in a black 96-well plate in 200 µL of complete media. MitoTracker^TM^ Deep Red (M22426, Invitrogen) was added to the wells at 500 nM and incubated for 30 minutes at 37 °C. Cells were washed with PBS and fixed using 4% (v/v) PFA for 15 minutes. After fixation, cells were washed, stained with Hoechst (2 µM), and imaged using ImageXpress Micro Confocal instrument. Fluorescence intensity of MitoTracker^TM^ was quantified using the Multi-Wavelength Cell Scoring Application Module for MetaXpress software and normalised per cell number (nuclei staining). Four images were acquired per well (n=2 wells per condition, N=3 independent experiments).

### Compound Profiling in Lysosomal Assays

Chloroquine (C6628, Sigma-Aldrich) and ammonium chloride (213330, Sigma-Aldrich) stocks were prepared in water at 50 mM and 5 M, respectively. Compounds were dispensed using ECHO Acoustic Liquid Handler (Beckman Coulter) to prepare a source plate containing stocks at 4x final assay concentration. From each stock, 10 μL were added per well to achieve the final assay concentration. 0.1% (v/v) water was included in each plate as a vehicle control. Data was normalised to the vehicle response.

### Statistical Analysis

Statistical analysis and graphical evaluations were carried out using Prism 10 (GraphPad). Data were plotted as violin plots to clearly represent the distribution and variability of the data. Comparisons between groups were performed using One-Way ANOVA followed by a *post hoc* test. A *P* < 0.05 was considered statistically significant. Descriptions of the statistical methods are provided in the legend of each figure.

## Results

We initially characterised lysosomal function in fibroblasts derived from healthy, AD and PD donors to understand whether these cells reflect lysosomal dysregulation reported in AD and PD. The fluorescence intensity of TMEM175 (a lysosomal expressed ion channel) and LAMP1 (glycoprotein located in lysosomes) was quantified in the presence of serum or under serum-starvation (**Figure 1**). Under serum culture conditions, PD fibroblasts showed reduced levels of TMEM175 (**Figure 1a,b**) with no differences in LAMP1 levels (**Figure 1c**). When serum-starved to induce autophagy and lysosomal activity, AD- and PD-derived fibroblasts continued to exhibit low levels of TMEM175 compared to the healthy donor (**Figure 1a,d**). However, a slight reduction in LAMP1 levels was found PD fibroblasts, in particular, for PD37 (**Figure 1e**). Given that similar trends were found for TMEM175 and LAMP1 at 4 (**Figure S1**) and 18 hours of serum-starvation (**Figure 1**), lysosome levels were further quantified using LysoTracker^TM^ at the later starvation time-point.

**Figure 1.**
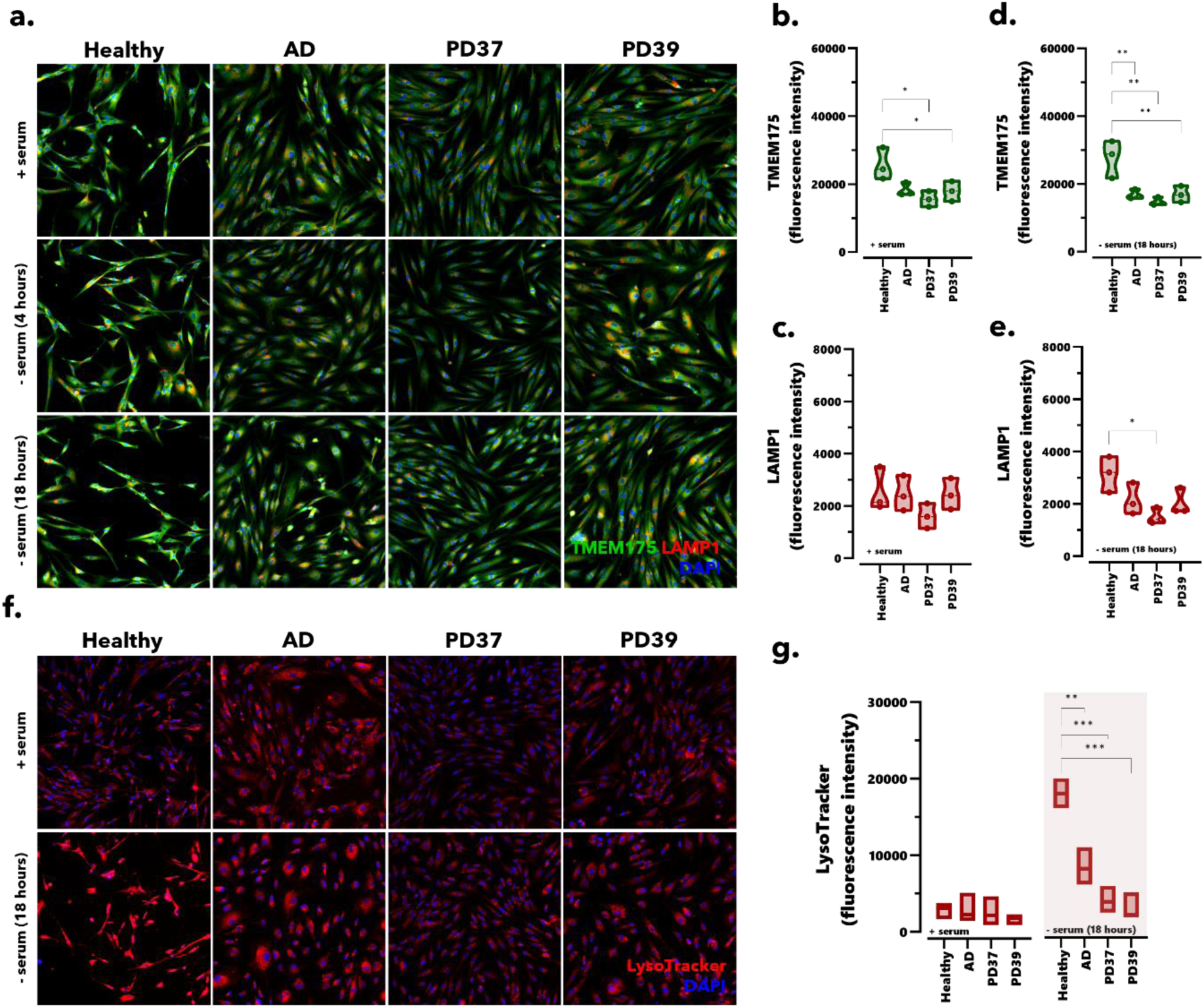
Fibroblasts derived from AD and PD donors exhibit altered lysosomal numbers when starved. (**A**) Representative images of TMEM175 (green) and LAMP1 (red) staining in human healthy, AD and PD-derived fibroblasts cultured in the presence of serum (+ serum) or under starvation (- serum) for 4 or 18 hours. The nuclei are represented in blue (DAPI). Images acquired at 20X magnification. Violin plots comparing the median levels of (**B**) TMEM175 and (**C**) LAMP1 cultured in the presence of serum. Violin plots comparing median (**D**) TMEM175 and (**E**) LAMP1 levels in fibroblasts starved for 18 hours. Fluorescence intensity normalised per cell number. (**F**) LysoTracker^TM^ staining (in red) in human fibroblasts from healthy, AD and PD donors under serum and starvation conditions for 18 hours. Nuclei counterstaining in blue. Images at 20X magnification. (**G**) Violin plots of LysoTracker^TM^ fluorescence intensity in fibroblasts in the presence and absence of serum (18 hours). Data represented as mean ± SD (N=3 independent experiments). * *P* < 0.05, ** *P* < 0.01, *** *P* < 0.001, One-Way ANOVA (Dunnett’s post-hoc) comparing healthy *versus* AD or PD fibroblasts.

LysoTracker^TM^ is a fluorescent probe that freely diffuses and accumulates in acidic organelles, such as lysosomes, enabling quantifying lysosomal levels in living cells. **Figure 1f** depicts the fluorescence of LysoTracker^TM^ Deep Red in fibroblasts cultured in serum or under serum starvation. In healthy and AD fibroblasts, serum-starvation triggered an increase in the LysoTracker^TM^ fluorescence intensity, which correlates with an increase in the number of lysosomes. Though the AD fibroblasts exhibited a lower dye accumulation level than the healthy donor. Both PD37 and PD39 fibroblasts exhibited similar levels of LysoTracker^TM^ accumulation in the presence and absence of serum (**Figure 1g**). These data indicate that under serum conditions, healthy, AD and PD fibroblasts present similar accumulation of LysoTracker^TM^. However, when serum-starved, AD and PD fibroblasts respond differently when compared to the healthy donor.

Given that the lysosomal levels in AD and PD fibroblasts under serum-starvation differed from those observed in the healthy donor, indicating changes in lysosomal function, we next investigated whether these fibroblasts exhibit any variations in lysosomal pH and enzymatic activity (**Figure 2**). Alterations in lysosomal pH were measured by incubating the fibroblasts cultured in either serum conditions or serum-starvation with LysoSensor^TM^ Yellow/Blue DND-160. Yellow:Blue fluorescence ratio of LysoSensor^TM^ dye gives indicates the acidity of lysosomes. An increase in this ratio signifies more acidic lysosomes. As shown in **Figure 2a**, no significant differences in lysosomal pH were found between healthy and AD, PD37 or PD39 fibroblasts, even when serum-starved for 18 hours. Activity of cathepsin B enzyme was also quantified in serum and serum-starvation culture conditions (**Figure 2b-d**). From the images of cathepsin B staining, serum-starvation appears to have no impact on enzymatic activity, except for fibroblasts derived from the AD donor that depicted increased fluorescence, indicating increased cathepsin B activity (**Figure 2b**). No statistically significant differences were found in the fluorescence intensity between healthy and AD or PD fibroblasts under serum conditions (**Figure 2c**); however, a significant increase in cathepsin B activity was observed for AD compared to healthy donor under serum-starvation (**Figure 2d**). Our data suggests that, as the lysosomal pH is unaltered compared to the healthy donor, cathepsin B enzyme activity remains functional in fibroblasts from AD and PD donors. Hence, this supports the impact of lysosomal pH in tightly regulating the activity of enzymes within the lysosome.

**Figure 2.**
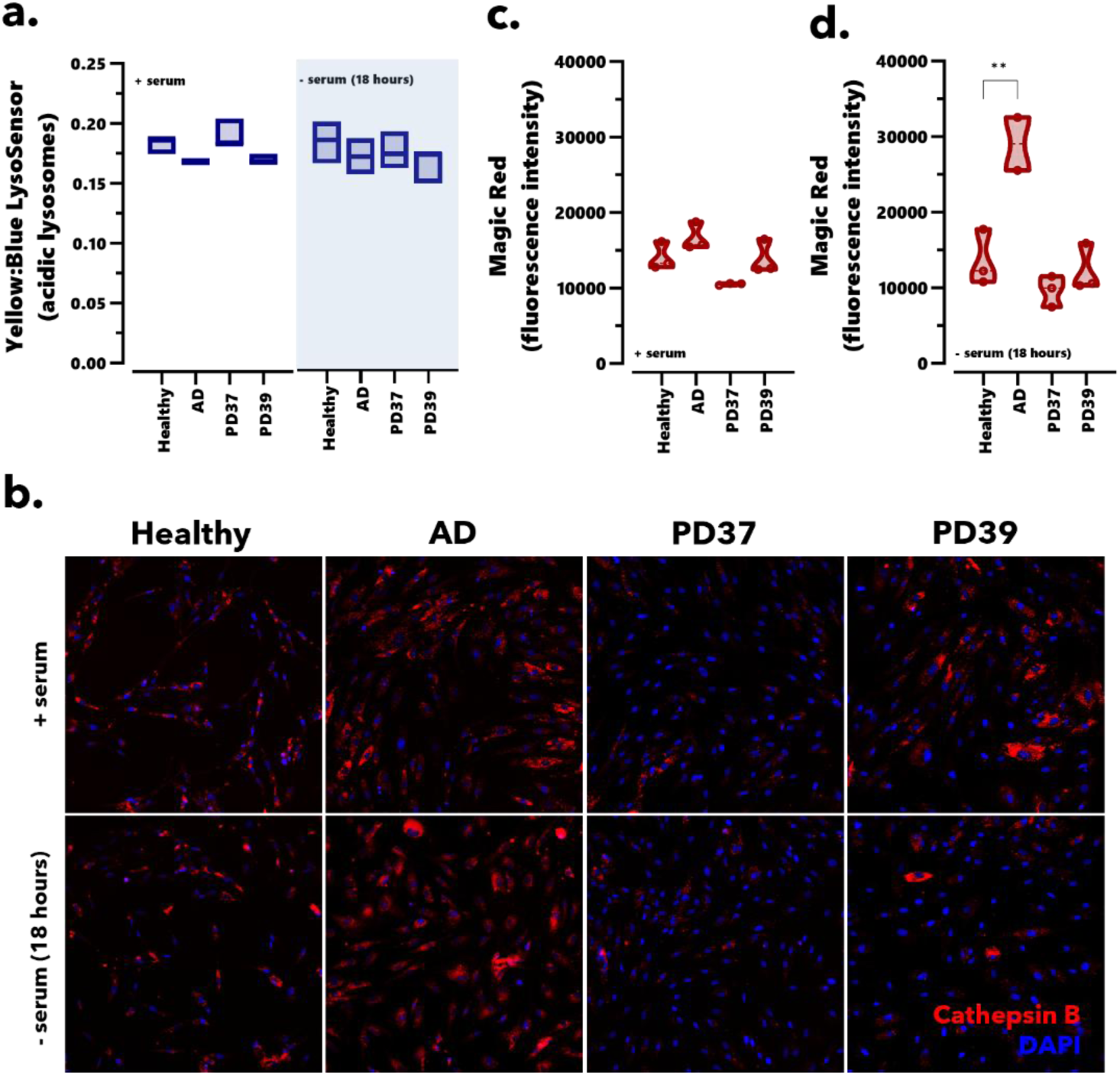
Fibroblasts derived from AD and PD donors present no alterations in lysosomal pH or cathepsin B enzyme activity compared to healthy donors. (**A**) Violin plots of LysoSensor^TM^ yellow:blue fluorescence ratio for human fibroblasts derived from healthy, AD and PD donors in the serum conditions and starvation for 18 hours. (**B**) Representative images of Magic Red cathepsin B staining in healthy, AD and PD fibroblasts cultured in the presence of serum (+ serum) or under starvation (- serum) for 18 hours. Nuclei are represented in blue (DAPI). Images acquired at 20X. Violin plots of Cathepsin B fluorescence intensity in fibroblasts cultured in the (**C**) presence of serum and (**D**) under starvation (18 hours). Fluorescence intensity was normalised per cell number. Data represented as the mean ± SD (N=3 independent experiments). ** *P* < 0.01, One-Way ANOVA (Dunnett’s post-hoc) comparing healthy *versus* AD.

We then evaluated autophagic flux in healthy, AD and PD fibroblasts by assessing p62 levels (**Figure S2**). Levels of p62 are used as an autophagic flux marker, with its degradation indicating effective autophagy. By binding to ubiquitinated substrates and LC3 on the autophagosome membrane, p62 facilitates the sequestration of damaged proteins and organelles into autophagosomes. Fibroblasts from AD and PD donors, cultured either in the presence of serum or under serum-starvation for 18 hours, showed significantly lower levels of p62 compared to the healthy donor (**Figure S2**). In serum conditions, AD and PD fibroblasts displayed statistically significant differences compared to healthy fibroblasts, suggesting greater basal autophagic degradation (**Figure S2b**). Serum-starving the fibroblasts appeared to have no impact on the autophagy flux of AD and PD fibroblasts, with PD fibroblasts continuing to present reduced levels of p62 compared to the healthy donor (**Figure S2c**). Based on the p62 levels, our data imply that AD and PD fibroblasts exhibit an altered autophagic flux, which may be connected to the lysosomal dysfunction found in these cells.

Considering the importance of the coordination between lysosomal and mitochondrial activity for healthy cellular metabolism ^5^, and increased evidence linking impaired mitochondrial function to PD pathology ^31,32^, we also explored potential changes in mitochondrial function in PD fibroblasts. As shown in **Figure 3a,b**, following FCCP stimulation in PD37 fibroblasts, a significant increase in Fluo-3 fluorescence was observed compared to the healthy donor. Our data indicates that mitochondria from PD37 fibroblasts were able to uptake and/or retain more mitochondrial calcium compared to healthy fibroblasts. Accordingly, FCCP increased the cytosolic fluorescence intensity of TMRE (a dye sequestered by active and negative mitochondria). Such findings suggest hyperpolarisation of the mitochondria and accumulation of TMRE in the matrix, which is released back to the cytosol upon FCCP stimulation (**Figure 3c,d**).

**Figure 3.**
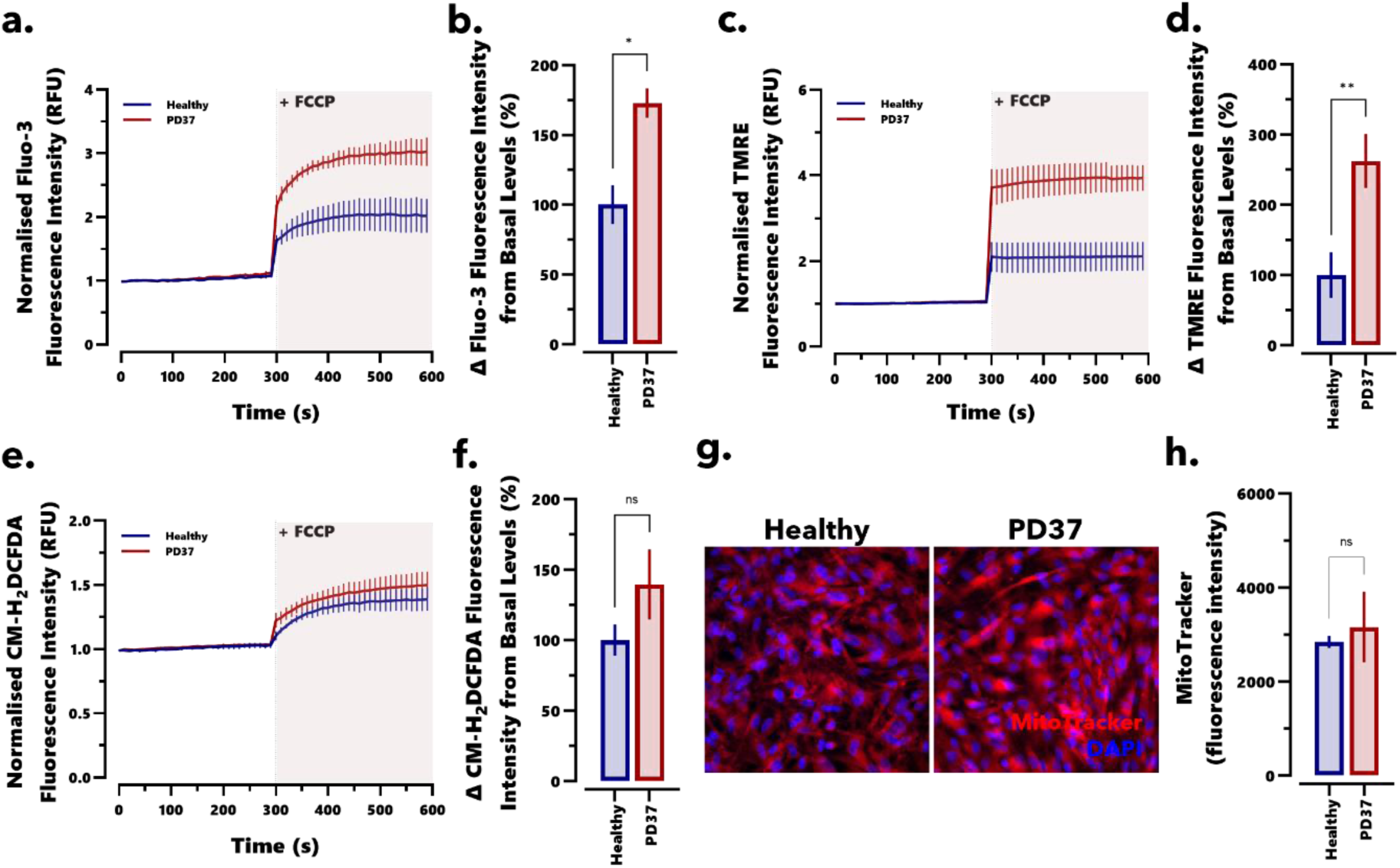
Fibroblasts from a PD donor exhibit altered mitochondrial dysfunction. (**A**) Fluo-3 fluorescence intensity, indicating intracellular calcium concentration, for healthy and PD37 fibroblasts. Fibroblasts were stimulated with FCCP following 5 minutes of baseline measurement, and readings were continued for another 5 minutes. Intensity readings were normalised to baseline recording. (**B**) Difference (Δ) in Fluo-3 fluorescence intensity pre- and post-FCCP stimulation. (**C**) TMRE fluorescence intensity (membrane potential indicator) in healthy and PD37 fibroblasts. (**D**) Difference (Δ) in TMRE fluorescence intensity pre- and post-FCCP stimulation. Data represented as mean ± SEM (N=4). (**E**) CM-H_2_DCFDA fluorescence intensity in healthy and PD37 fibroblasts. (**F**) Change in CM-H2DCFDA fluorescence intensity pre- and post-FCCP stimulation. Mean ± SEM (N=5 independent experiments). (**G**) Representative images of MitoTracker^TM^ staining in healthy and PD37 fibroblasts. Images were acquired at 40X. Nuclei are shown in blue (DAPI). (**H**) MitoTracker^TM^ fluorescence intensity quantification in healthy and PD fibroblasts. Mean ± SD (N=3 independent experiments). * *P* < 0.05, ** *P* < 0.01, Student’s t-test comparing healthy and PD37 fibroblasts.

Since mitochondrial calcium and membrane potential are associated with oxidative stress levels, we further investigated ROS levels in the fibroblasts. As illustrated in **Figure 3e**, a minor increase in CM-H_2_DCFDA fluorescence levels was found in PD37 fibroblasts, after addition of FCCP compared to healthy fibroblasts (**Figure 3f**), although not statistically significant. Such findings suggest that ROS production and oxidative stress were not affected in the PD37 fibroblasts, even in the presence of calcium handling and membrane potential alterations.

Given that the increase in cytosolic calcium and TMRE following FCCP stimulation may indicate a rise in mitochondrial content, we examined the total number of mitochondria in healthy and PD37 fibroblasts using MitoTracker^TM^ staining (**Figure 3g**). No statistically significant differences were obtained between healthy and PD donors (**Figure 3h**). However, MitoTracker^TM^ assay may lack the sensitivity to fully explain the observed mitochondrial hyperpolarisation or to understand other unrelated mechanisms that contributed to such mitochondrial dysfunction.

Together, these findings establish that patient-derived fibroblasts exhibit marked alterations in lysosomal function, including changes in lysosome abundance and autophagic degradation capacity, as well as impairments in mitochondrial function. Importantly, the assays developed to evaluate lysosomal activity in these fibroblasts are based on high-throughput imaging, making them highly suitable for compound screening in drug discovery. Thus, these results highlight the value of using patient-derived fibroblasts as a robust cellular model for identifying and evaluating modulators of lysosomal function and related pathways.

To explore this potential, we next selected a set of established imaging-based assays, including LAMP1, LysoTracker^TM^ and Magic Red Cathepsin B, to profile compounds known to modulate lysosomal function, CQ and AC ^33,34^. CQ and AC are weak bases that diffuse across cellular membranes in their uncharged forms. Upon entering acidic lysosomes, both CQ and AC become protonated and accumulate in the lysosomes due to the pH gradient; this leads to lysosomal alkalinisation, thereby impairing the activity of acid-dependent hydrolases and disrupting lysosomal function.

To establish the optimal incubation time with the two compounds, we conducted time-course experiments by incubating healthy and PD37 fibroblasts with CQ and AC for 1 to 24 hours (**Figure S3**). LAMP1 and LysoTracker^TM^ assays found an effect with more extended incubation periods, while the Magic Red Cathepsin B assay detected alterations in enzymatic activity within 1-2 hours. CQ induced an increase in LAMP1, LysoTracker^TM^, and Magic Red signal, which reflects an increased level of lysosomes (**Figure 4**); however, no statistical significance was found for AC in LAMP1 and LysoTracker^TM^ assays (**Figure 4**). A response was only obtained for AC in the Magic Red Cathepsin B assay after a more prolonged incubation time (i.e., 24 hours). Interestingly, healthy fibroblasts appear more responsive to CQ than the PD37 fibroblasts. EC_50_ values of 0.83 and 2.52 µM were obtained for CQ in a Magic Red cathepsin B assay.

**Figure 4.**
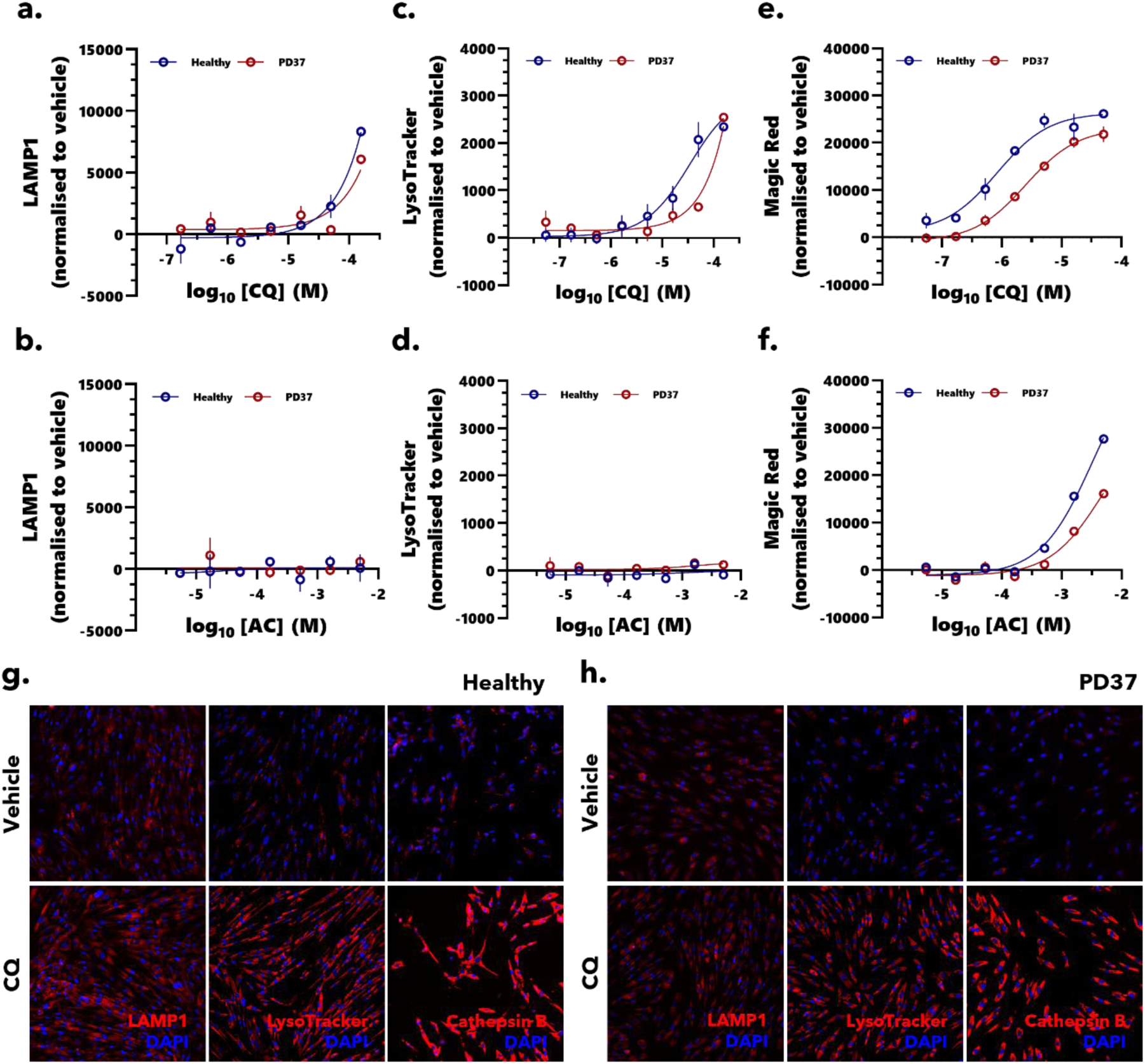
Profiling of CQ and AC in both healthy and PD-derived fibroblasts across different lysosomal assays. Concentration-response curves for CQ and AC in healthy and PD37 fibroblasts in (**A-B**) LAMP1, (**C-D**) LysoTracker^TM^, and (**E-F**) Magic Red cathepsin B assays. Fluorescence intensity was normalised to the vehicle (0.1% v/v water). Data represented as mean ± SD (N=3 independent experiments). Representative images of LAMP1, LysoTracker^TM^ and Magic Red cathepsin B staining in healthy (**G**) and PD37 (**H**) fibroblasts treated with CQ (at 150 µM) or vehicle (0.1% v/v water) for 24 hours (or 4 hours for the Magic Red assay). Images acquired at 20X.

Our findings reveal CQ as a valuable compound to validate the robustness and sensitivity of the developed assays, demonstrating their reliability in detecting lysosomal alterations and confirming their suitability for compound screening in drug discovery workflows.

## Discussion

Given the essential role of lysosomal and mitochondrial function in maintaining cellular homeostasis, and their involvement in neurodegenerative diseases, we investigated whether patient-derived fibroblasts recapitulate key dysfunctions reported in AD and PD. In addition, we assessed the suitability of these patient-derived fibroblasts as a platform for profiling therapeutic compounds aimed at restoring lysosomal function in the context of neurodegeneration.

Lysosomal dysfunction has been increasingly recognised as a main contributor to the pathogenesis of both AD and PD ^26^. To investigate this further, we assessed several parameters of lysosomal function in patient-derived fibroblasts, including lysosomal abundance, pH levels, enzymatic activity, and autophagic flux. We first examined the levels of TMEM175 and LAMP1, two lysosomal markers, and found statistically significant differences in fibroblasts from AD and PD patients compared to healthy controls, particularly under serum-starvation conditions.

TMEM175, in particular, has been directly linked to PD risk ^35^. Loss or dysfunction of TMEM175 leads to lysosomal over-acidification, reduced cathepsin B and glucocerebrosidase activity, and impaired autophagy and mitochondrial function ^36,37^. Two common TMEM175 variants have been identified: a risk-associated p.M393T (rs34311866) variant and a protective p.Q65P (rs34884217). p.M393T mutation causes a significant decrease in the stability of TMEM175 structural domain, damaging protein transport and maturation, which results in reduced localisation of TMEM175 in the lysosome. Carriers of this mutation exhibit pathological features similar to a TMEM175 knockout (KO), with a dysregulated lysosomal pH, defective clearance of autophagy substrates and increased accumulation of α-syn ^36^. Unlike the loss-of-function pM393T, p.Q65P is the protective variant, as it stabilises the ion-selective section of the structural domain, and enhances permeability of the TMEM175 channel ^38^. Although the presence of specific TMEM175 mutations was not assessed in the PD fibroblasts, the decrease in levels observed in both PD lines may reflect functional impairment of TMEM175, with potential consequences for lysosomal activity.

In terms of LAMP1 levels, no significant differences were observed between fibroblasts from AD or PD patients and those from the healthy donor under serum-supplemented conditions. These results suggest that lysosomal abundance remains unchanged in patient-derived fibroblasts under these conditions. This observation aligns with previous studies reporting no alterations in LAMP1 levels in idiopathic and glucocerebrosidase-mutant PD-derived fibroblasts ^8,39,40^. Interestingly, under nutrient-deprived conditions, PD37 fibroblasts displayed a reduction in lysosomal levels. This may reflect a reduced ability to cope with stress and impaired capacity to clear cellular components upon autophagy induction.

To further investigate lysosomal abundance and activity, we performed LysoTracker™ staining. When cultured in serum-supplemented media, no significant differences between fibroblasts from healthy donors and AD or PD patients were observed. These findings are consistent with previously published data showing no alterations in LysoTracker™ accumulation in PD versus healthy fibroblasts ^40^. However, under serum-starvation, healthy fibroblasts exhibited a statistically significant increase in LysoTracker™ accumulation, indicating an adaptive increase in lysosomal numbers and/or activity. In contrast, PD fibroblasts failed to mount a similar response, further supporting the hypothesis of an impaired lysosomal adaptation to stress in these cells.

Although LAMP1 is widely recognised as a lysosomal marker, it may not always provide a sensitive readout for lysosomal deficits ^41^. Therefore, confirming lysosomal levels using complementary approaches, such as LysoTracker™, is crucial. Taken together, these findings suggest that fibroblasts derived from AD and PD patients display a limited response to stress conditions, such as nutrient starvation, which is reflected by a reduced ability to increase lysosomal levels and activity.

Following the indication of defective lysosomal activity, we next evaluated lysosomal pH and enzymatic function in patient-derived fibroblasts. Assessment of lysosomal pH using the LysoSensor^TM^ assay revealed no significant differences between fibroblasts from healthy donors and those from AD or PD patients, suggesting that lysosomal acidity is maintained across all cell lines. Coffey *et al.* ^42^ reported that AD-derived fibroblasts with a PSEN1 mutation (A264E) exhibit elevated lysosomal pH compared to a healthy donor; however, evidence on lysosomal pH in human fibroblasts from AD and PD donors is limited.

We then explored lysosomal enzymatic activity by measuring cathepsin B activity. Interestingly, under serum-starvation conditions, AD fibroblasts displayed a statistically significant increase in cathepsin B activity compared to the healthy donor. This observation is consistent with several studies reporting elevated cathepsin B protein levels or activity in plasma and cerebrospinal fluid from AD patients, and amyloid plaques ^43–45^. Genetic investigation suggests that cathepsin B exerts a protective effect against PD ^46^. However, in fibroblasts derived from a PD donor, no alterations were described in cathepsin B levels or activity ^47^, which is consistent with our data.

Together, these findings suggest that the increased LysoTracker™ accumulation observed in some fibroblasts is likely due to an increase in lysosomal number rather than alterations in lysosomal acidity, as supported by the LysoSensor™ assay results.

In light of the pivotal role of autophagy dysfunction in the pathogenesis of neurodegenerative diseases ^48^, we next investigated p62 levels - a key marker of autophagic activity. Interestingly, AD and PD fibroblasts displayed consistently reduced p62 levels relative to the healthy donor, suggesting potential alterations in autophagy regulation or protein clearance mechanisms. Other reports described comparable findings in different fibroblasts from sporadic and juvenile PD donors, and donors with specific mutations, such as LRRK2 and PARKIN mutations ^21,40,49^. Accordingly, it has been implied that this alteration in p62 might occur as an auto-activated compensatory mechanism of degradation, which is challenging to sustain and leads to a buildup of toxic protein aggregates ^50^.

Given the well-established association between mitochondrial dysfunction and PD, we investigated the mitochondrial function in patient-derived fibroblasts. In our study, PD37 fibroblasts showed alterations in both mitochondrial calcium uptake/retention and mitochondrial membrane potential. Specifically, we observed increased mitochondrial calcium levels and signs of hyperpolarisation. Accumulation of calcium in mitochondria, which leads to calcium overload, has been associated to PINK1 deficiency, as found in autosomal recessive PD ^51^. Additionally, PINK1 deletion-induced mitochondrial dysfunction has been shown to impair lysosomal activity ^52^, evidencing the significance of mitochondrial and lysosomal crosstalk in PD. Interestingly, while Mortiboys *et al*. ^53^ reported a decrease in mitochondrial membrane potential in fibroblasts from early-stage PD patients and those carrying PARKIN mutations, other studies in idiopathic PD-derived fibroblasts have shown the opposite, an increase in membrane potential compared to healthy controls ^54^. These findings align with our results and highlight that the mechanism of mitochondrial dysfunction in PD may vary depending on disease aetiology.

Mitochondrial dysfunction, often occurring alongside impaired autophagy, has previously been reported in fibroblasts derived from PD patients ^49,55^. In line with these findings, the mitochondrial alterations observed in PD37 fibroblasts, including increased mitochondrial calcium levels and hyperpolarisation, may contribute to the lysosomal dysfunction and reduced p62 levels also identified in these cells. It is possible that defective mitochondria in PD37 fibroblasts trigger a compensatory response, activating autophagy and lysosomal pathways in an attempt to promote mitochondrial clearance and maintain cellular homeostasis.

Following our evaluation of lysosomal and mitochondrial function in patient-derived fibroblasts, we proceeded to validate lysosomal assays for compound profiling in a drug discovery setting. A range of compounds, including CQ and AC, produced distinct effects on lysosomal function, which were detected using our established suite of functional assays. Previous studies have demonstrated that CQ, by diffusing into lysosomes and becoming protonated, raises lysosomal pH and impairs lysosomal function ^33,56,57^. In our study, CQ treatment led to increased LAMP1 expression and LysoTracker™ accumulation, along with elevated Cathepsin B activity in healthy and PD37 fibroblasts. These findings are consistent with lysosomal accumulation due to impaired autophagosome–lysosome fusion and degradation ^57^.

Although AC has also been reported to inhibit lysosomal function by alkalinizing the lysosomal lumen ^34; 58^, its effects in our study appeared limited to lysosomal enzyme activity, as detected by the Magic Red Cathepsin B assay, with no significant impact on other lysosomal readouts. A plausible explanation is that AC, while acting as a weak base, does not accumulate in lysosomes to the same extent as other lysosomotropic agents such as CQ. Its alkalinizing effects are typically more transient and less potent ^59^, which may suffice to reduce the activity of pH-sensitive lysosomal enzymes, but not to induce broader structural or functional disruptions detectable by other assays (e.g., LysoTracker™ staining, LAMP1 expression). In contrast, CQ is an amphiphilic molecule that is retained within lysosomes, resulting in sustained pH elevation and a more comprehensive disruption of lysosomal function, including inhibition of autophagosome–lysosome fusion ^33^. Thus, the limited response observed with AC may reflect its distinct pharmacokinetic properties and weaker lysosomotropic profile compared to CQ.

To further strengthen the translational value of this drug discovery platform, future work should focus on validating these lysosomal assays using compounds that specifically target lysosome-associated proteins. One promising example is TMEM175, a lysosomal potassium channel involved in maintaining lysosomal membrane potential and pH homeostasis, and genetically linked to Parkinson’s disease. By assessing the effects of TMEM175 modulators and other lysosome-targeted compounds, we aim to refine the sensitivity and specificity of the assay suite, enhancing its application in high-throughput screening for novel therapeutics in neurodegeneration.

In summary, we extensively characterised lysosomal and mitochondrial function in patient-derived fibroblasts from individuals with AD and PD, highlighting cellular dysfunctions relevant to disease pathology. Importantly, we demonstrated that several assays are robust, sensitive, and compatible with the profiling of compounds in lysosome-relevant assays. These findings not only validate human patient-derived fibroblasts as a relevant *in vitro* cellular model but also support their application in broader drug discovery workflows, enabling the identification and profiling of novel compounds targeting lysosomal and mitochondrial pathways in the context of neurodegeneration.

## Acknowledgements

We gratefully acknowledge the Coriell Institute for providing the human fibroblasts used in this study. The following cells were obtained from the NIA Aging Cell Culture Repository: AG02262, AG06840, AG20446, and AG20442. We also acknowledge Foteini-Nafsika Damaskinaki for her assistance during the initial phases of the mitochondrial assays and cell culture work.

## Funding

The authors received no financial support for the research, authorship, and publication of this article.

## Authors Contributions

DL, RL and PM carried out experiments. DL, PM, and TR wrote the manuscript. DL designed the study, wrote the protocols, and revised the manuscript. DL and TR wrote the final version. All authors approved the final version of the manuscript.

## Data Availability

All the results obtained or analysed throughout our study are presented in this manuscript (and its Supplementary Information file).

## Competing Interests

The authors declare the following competing financial interest: Diana M. Leite, Philippa Malko, Stuart Thomson, Tatiana R. Rosenstock, Tim Philipps, and Naheed R. Mirza were full-time employees of Sygnature Discovery Limited at the time the work was completed.

