## Supplementary material for "Profiling Lysosomal and Mitochondrial Dysfunction in Neurodegenerative Diseases Using Human Fibroblasts for Translational Therapeutic Screening": Figure S1, Figure S2, Figure S3

[1] Sygnature Discovery, BioCity, Nottingham, United Kingdom.

[2] Nottingham University, Nottingham, United Kingdom.

*** Corresponding Author:**

Diana M. Leite, Tatiana R. Rosenstock

Sygnature Discovery, BioCity,

Pennyfoot Street, Nottingham, United Kingdom

, t.rosenstock@sygnaturediscovery

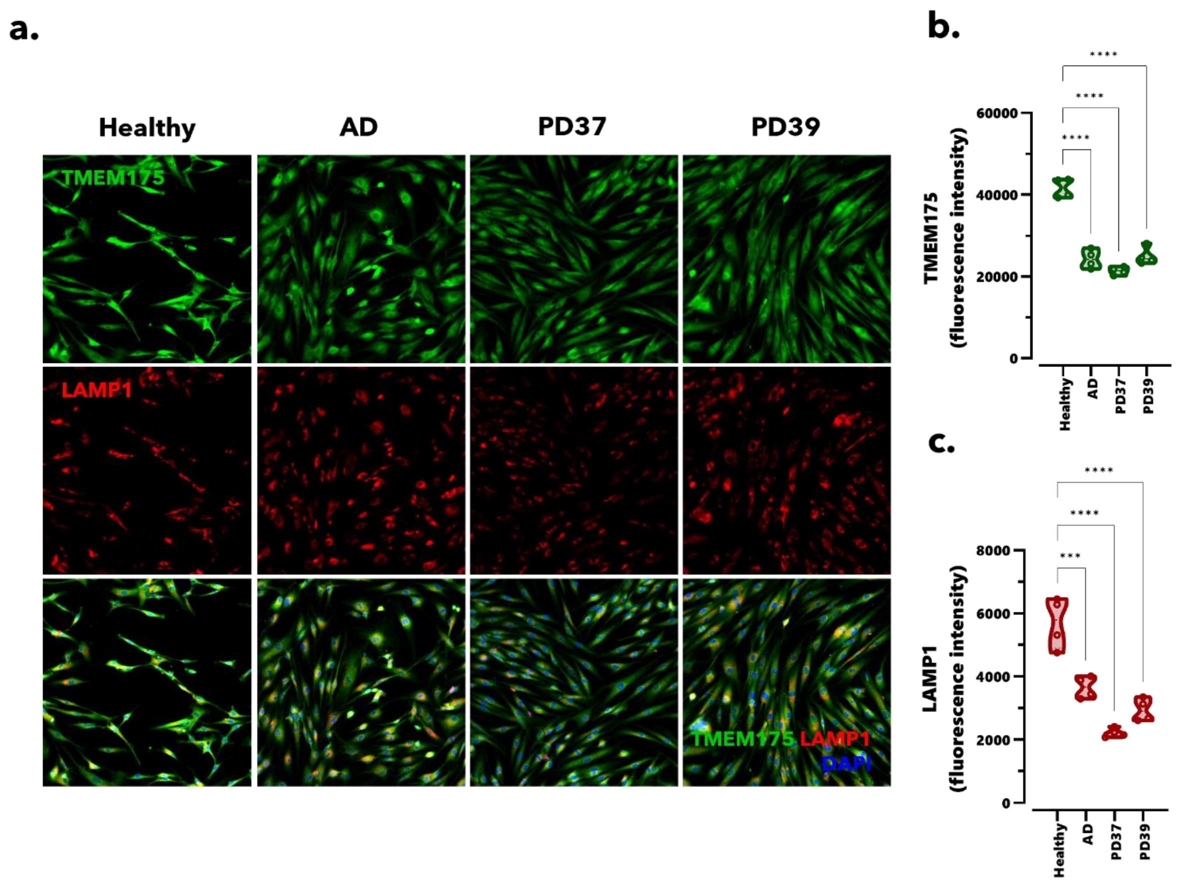


**Supplementary Figure 1. Fibroblasts derived from AD and PD donors exhibit altered lysosomal numbers when serum-starved for 4 hours.**

(**A**) Representative images of TMEM175 (green) and LAMP1 (red) staining in human healthy, AD and PD-derived fibroblasts cultured under serum-starvation for 4 hours. Nuclei are shown in blue. Images acquired at 20X magnification. Violin plots comparing the median levels of (**B**) TMEM175 and (**C**) LAMP1 in fibroblasts starved for 4 hours. Fluorescence intensity normalised per cell number. Data represented as mean ± SD (N=4 independent experiments, 4 wells per condition in each experiment). *** $P < 0.001$, **** $P < 0.0001$, One-Way ANOVA (Dunnett's post-hoc) comparing healthy *versus* AD and PD.

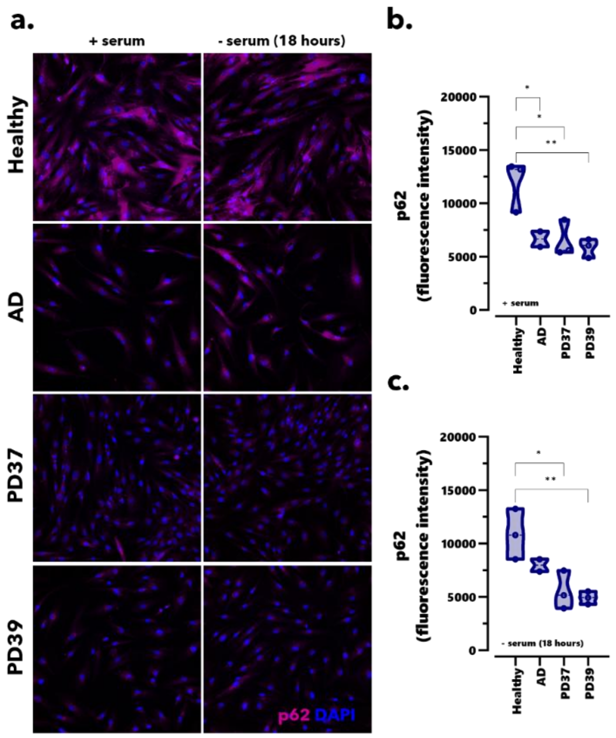


**Supplementary Figure 2. Fibroblasts from AD and PD donors display altered autophagic flux.** (**A**) p62 staining in healthy, AD, and PD-derived fibroblasts maintained in serum or under serum-starvation (18 hours). Nuclei are shown in blue and p62 in magenta. Images acquired at 20X magnification. Violin plots of the p62 fluorescence intensity quantification in fibroblasts (**B**) in serum-supplemented and (**C**) in starvation conditions. * $P < 0.05$, ** $P < 0.01$, One-Way ANOVA (Dunnett's post-hoc) comparing healthy *versus* AD and PD fibroblasts (N=3 independent experiments, 4 wells per condition in each experiment).

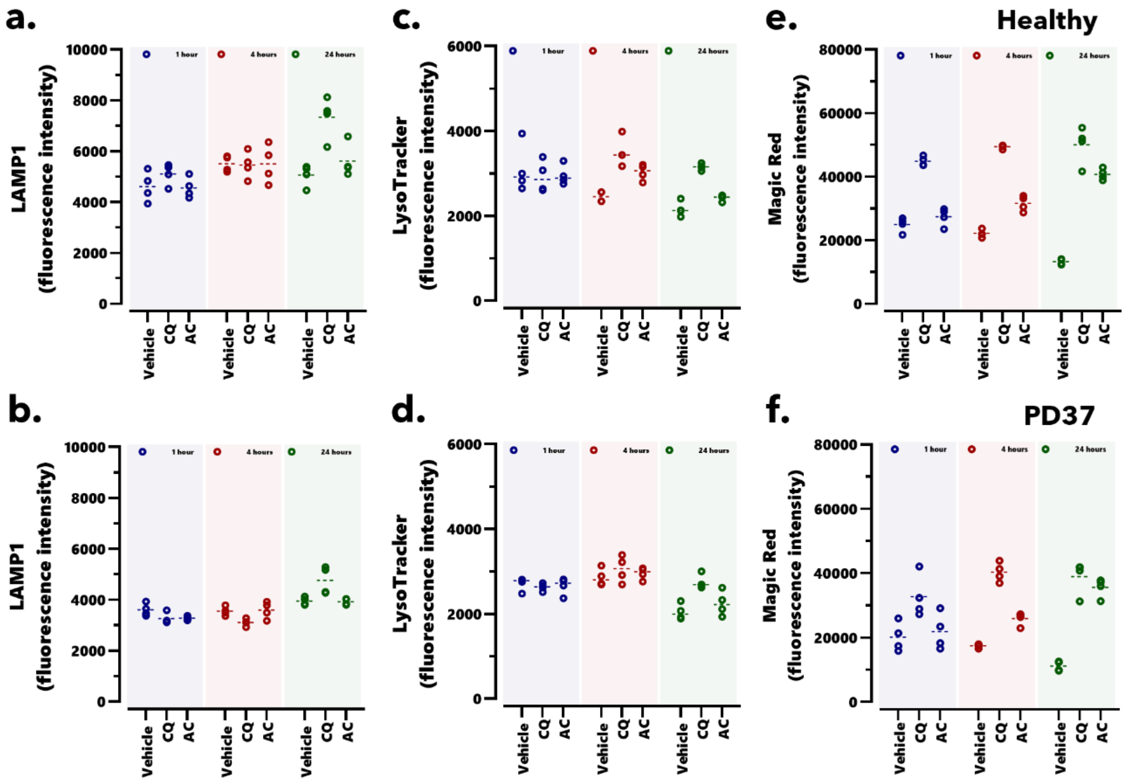


**Supplementary Figure 3. Time-course of CQ and AC stimulation in healthy and PD37 fibroblasts**

Profiling of CQ (50 μM) and AC (5 mM) in LAMP1 (**A,B**), LysoTracker™ (**C,D**) and Magic Red (**E,F**) assays in time-course using healthy (top panel) and PD37 (bottom panel) fibroblasts. Fluorescence intensity normalised to cell number. Data represented as mean ± SD (n=4 wells).
